# Optogenetic silencing of primary afferents reduces evoked and ongoing bladder pain

**DOI:** 10.1101/167841

**Authors:** Vijay K. Samineni, Aaron D. Mickle, Jangyeol Yoon, Jose G. Grajales-Reyes, Melanie Pullen, Kaitlyn Crawford, Kyung Nim Noh, Graydon B. Gereau, Sherri Vogt, H. Henry Lai, John A. Rogers, Robert W. Gereau

## Abstract

Patients with interstitial cystitis/bladder pain syndrome (IC/BPS) suffer from chronic pain that severely affects quality of life. Although the underlying pathophysiology is not well understood, inhibition of bladder sensory afferents temporarily relieves pain. Here, we explored the possibility that optogenetic inhibition of bladder sensory afferents could be used to modulate bladder pain. Specifically, we chose to study the role of Na_v_1.8^+^ sensory afferents before and after induction of a mouse model of bladder pain. The light-activated inhibitory proton pump Archaerhodopsin (Arch) was expressed under control of the Na_v_1.8^+^ promoter to selectively silence these neurons. Optically silencing Na_v_1.8^+^ afferents significantly blunted the evoked visceromotor response to bladder distension and led to small but significant changes in bladder function. To study of the role of these fibers in freely behaving mice, we developed a fully implantable, flexible, wirelessly powered optoelectronic system for the long-term manipulation of bladder afferent expressed opsins. We found that optogenetic inhibition of Na_v_1.8^+^ fibers reduced both ongoing pain and evoked cutaneous hypersensitivity in the context of cystitis, but had no effect in uninjured, naïve mice. These results suggest that selective optogenetic silencing of bladder afferents may represent a potential future therapeutic strategy for the treatment of bladder pain.

## Introduction

Interstitial cystitis/bladder pain syndrome (IC/BPS) is a debilitating pelvic pain syndrome of unknown etiology ^1,2^. IC/BPS patients experience chronic pelvic pain symptoms, including pain on bladder filling, increased urinary urgency and frequency as the primary clinical symptoms ^3-12^. Blocking afferent drive from the bladder by instillation of local anesthetics has long been known to reduce bladder pain in many patients with IC/BPS ^13,14^. More targeted desensitization of C-fiber afferents using resiniferatoxin (a potent TRPV1 agonist) has also been demonstrated to reduce bladder pain in IC/BPS patients, suggesting that targeted strategies to silence bladder nociceptive afferents hold promise for the clinical management of IC/BPS ^15,16^.

The TTX-resistant sodium channel Na_v_1.8 is expressed in nociceptor populations that respond predominantly to capsaicin and inflammatory mediators ^17-19^. Previous studies have shown the expression of Na_v_1.8 in C-fiber afferent neurons that innervate the bladder ^20,21^. Additionally, C-fiber bladder afferent neurons that express Na_v_1.8 exhibit increased excitability after induction of bladder cystitis ^22^. Recent studies demonstrate that rodents in which Na_v_1.8 gene is deleted or suppressed, exhibit diminished bladder nociceptive responses and referred hyperalgesia after induction of cystitis ^23,24^. These results suggest that selective silencing of primary afferent neurons expressing Na_v_1.8 could be an effective strategy for treating inflammation-induced bladder pain. However, the major caveat with these knockdown studies is that it is hard to control for compensatory upregulation of other Na_v_ channels. To overcome this obstacle, we took advantage of optogenetic techniques that can specifically and reversibly inhibit this specific population of neurons.

Archaerhodopsin (Arch), is a light-activated proton pump, which upon activation by green light (520-590 nm), induces membrane hyperpolarization and robustly suppresses neuronal firing ^25,26^. Previous studies have shown that transdermal illumination of Arch expressed in Na_v_1.8^+^ peripheral nerve terminals innervating the paw results in a significant decrease in somatic pain, suggesting that optogenetics can be used to effectively silence the peripheral nociceptors during chronic pain ^27,28^. Whether this approach can be extended to modulation of visceral pain is not known. Here, we test the hypothesis that Arch expression in Na_v_1.8^+^ sensory neurons can reduce bladder pain in the cyclophosphamide (CYP)–induced cystitis model of bladder pain in mice.

A major challenge in applying optogenetics to studies of viscera is achieving robust and consistent light delivery to the end organ in awake and behaving animals. Recent advances in wireless optoelectronics have made it possible to activate opsins expressed in the brain, spinal cord and peripheral neurons in freely moving animals ^29-31^. Here, we have developed ultra-miniaturized, optoelectronic devices that contain micro-scale light emitting diodes (μ-ILEDs) capable of wireless operation through near field communication (NFC) hardware. This device is designed to interface with peripheral abdominal tissue to illuminate the optogenetic probes expressed in the bladder afferents of freely moving mice. In this study, we utilize these new devices to investigate whether optical silencing of Na_v_1.8^+^ bladder sensory neurons can attenuate ongoing pain and referred visceral hyperalgesia associated with development of CYP-induced cystitis in freely moving animals.

## Results

### Immunohistochemical and electrophysiological characterization of Na_v_1.8-Arch expressing bladder sensory afferents

We crossed heterozygous Na_v_1.8-cre mice ^32^ to homozygous Ai35 mice ^33^ to generate Na_v_1.8-Arch mice. The retrograde tracer cholera toxin subunit B (CTB) was injected into the bladder wall of Na_v_1.8-Arch mice to identify Na_v_1.8-Arch-GFP^+^ neurons that project to the bladder. Seven days after CTB injection, dorsal root ganglia (DRG) at L6-S1 levels exhibited numerous CTB^+^ neurons (Fig. 1A-C). Quantitative analysis revealed that 75.2 ± 5.2% of CTB^+^ bladder projecting DRG neurons co-label with Arch-GFP. Amongst the CTB^+^ bladder-projecting DRG neurons that express Na_v_1.8-Arch-GFP, 42 ± 10.9% are NF200-positive (Fig. 1A), 41± 0.1% are CGRP-positive (Fig. 1B) and 32 ± 3.1% are IB4-positive (Fig. 1C). Whole mount staining of the bladder wall of Na_v_1.8-Arch mice showed that Arch-GFP-positive fibers co-label with the neuronal marker β3-tubulin, suggesting that Arch-GFP is effectively transported to afferent endings in the bladder (Fig. 1D).

**Figure 1.**
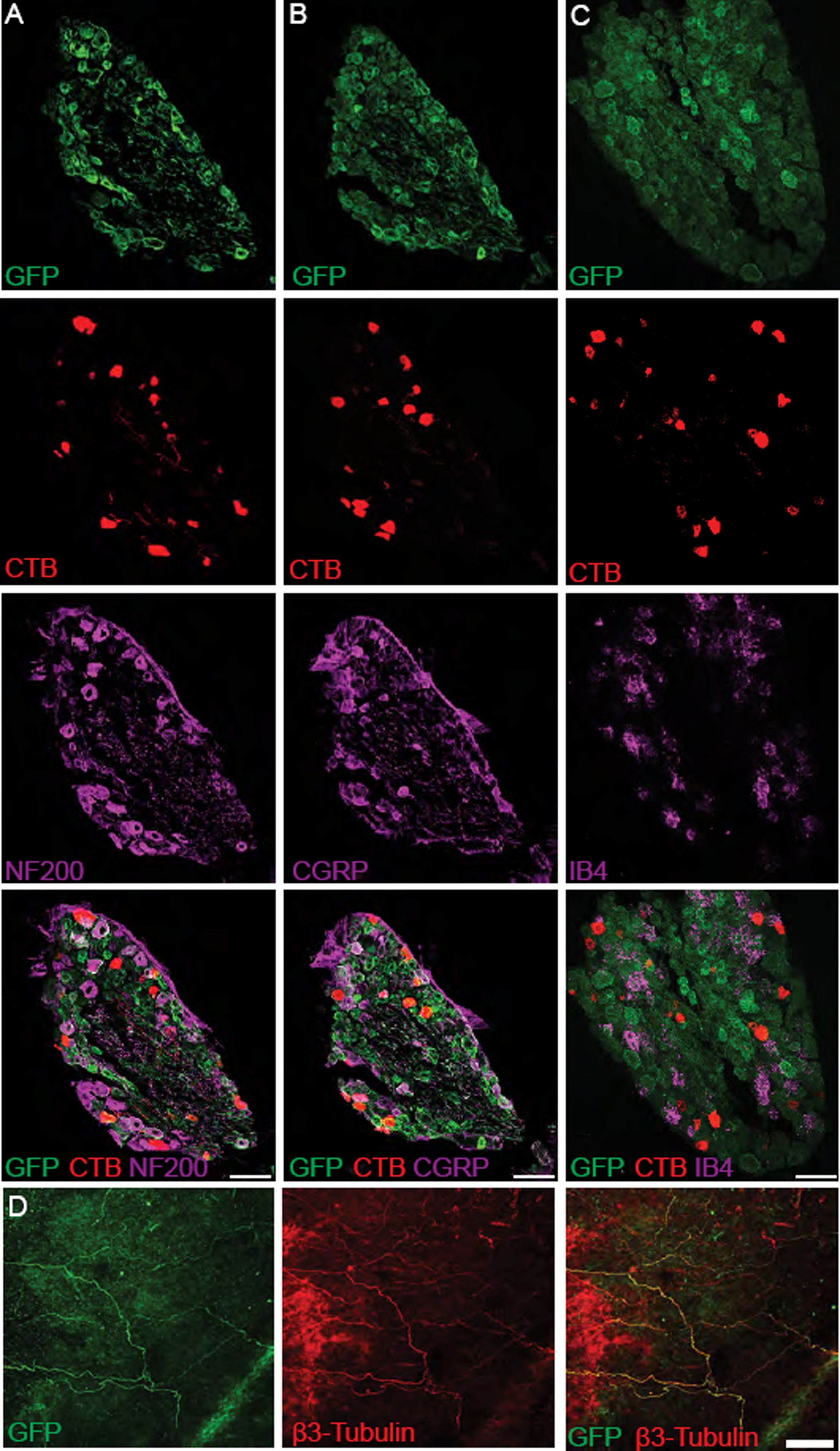
Histological characterization of L6-S1 dorsal root ganglion (DRG) neurons that project to the bladder from Na_v_1.8-Arch mice. (A, B & C) Immunohistochemical analysis of tissue from Na_v_1.8-Arch mice confirms eGFP reporter expression in L6-S1 DRGs. Labeling of bladder-innervating DRG neurons was achieved via bladder injections of the retrograde neuronal tracer cholera toxin B. Co-labeling of eGFP and CTB-555 with the neuronal markers calcitonin gene related peptide (CGRP), neurofilament 200 (NF200) and isolectin B4 (IB4) reveal a mixed population of sensory neurons that innervate the mouse bladder; scale bar 100 μm (n=2). (D) Whole mount staining of the bladder wall shows sensory neurons expressing eGFP innervate the bladder wall of Na_v_1.8-Arch mice and co-label with β3-tubulin; scale bar 100 μm.

We performed whole-cell patch clamp electrophysiology recordings on cultured DRG neurons from Na_v_1.8-Arch mice to verify functional expression of Arch in Na_v_1.8^+^ bladder sensory afferents. We identified bladder-projecting neurons, using the retrograde dye, DiI, injected into the bladder wall and then recorded from bladder-projecting (DiI^+^, red) and Arch-GFP^+^ (green) cell bodies (Fig 2A). In voltage clamp, illumination of Arch-GFP-expressing bladder-projecting neurons with green light (530 nm) induced robust outward currents (Fig. 2B). This current was sufficient to suppress action potential firing in response to ramp current injections in all neurons tested (Fig. 2C-D, *** p=0.001. n=7 neurons; Paired t test).

**Figure 2.**
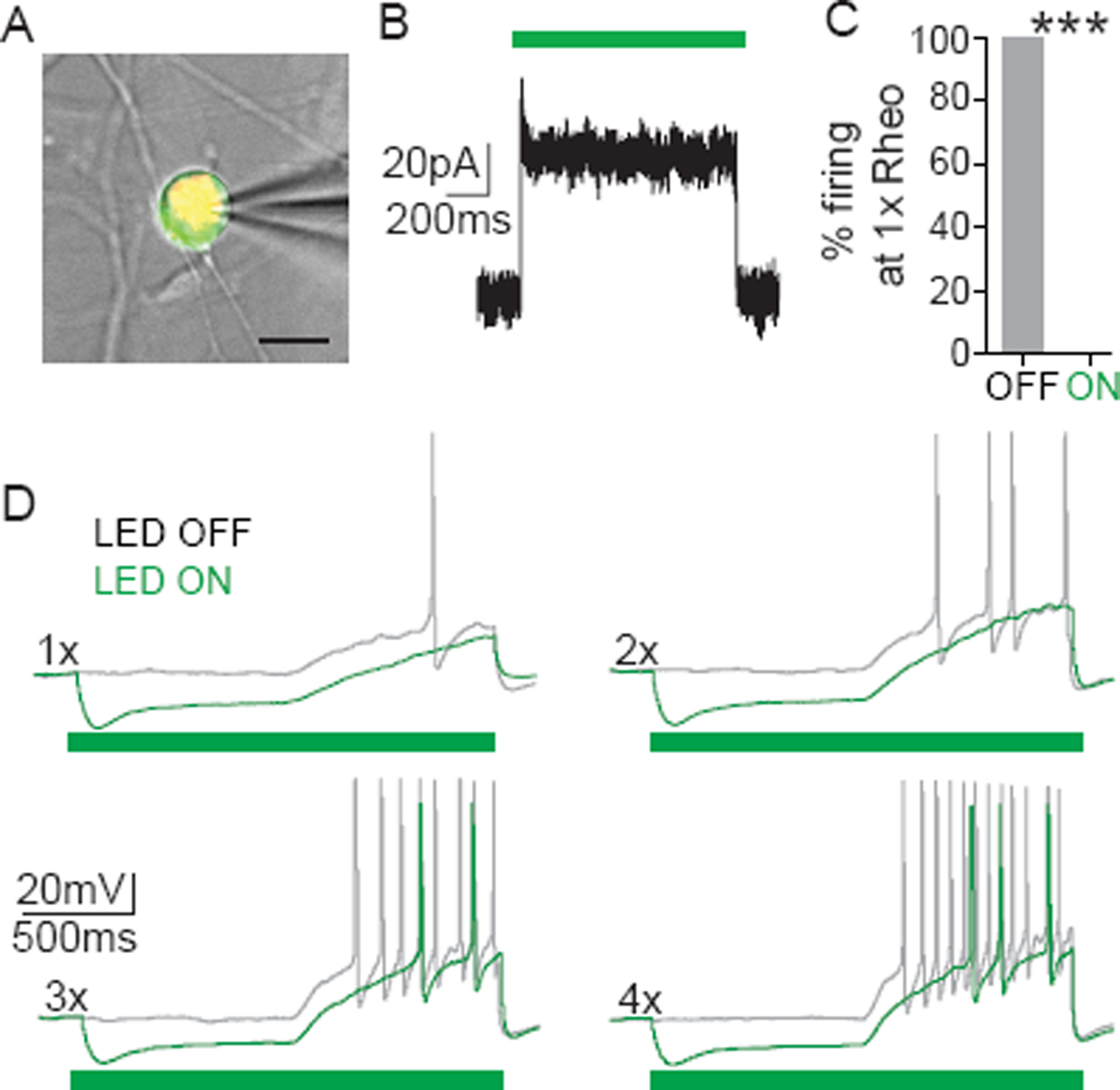
Arch activation in bladder sensory neurons decreased neuronal excitability. (A) Example of a patched neuron (21 μm diameter); green fluorescence indicates Arch-GFP expression, red fluorescence is DiI labeling, indicating that the neuron projects to the bladder wall and expressed Arch (yellow); scale bar = 20 μm. (B) Representative voltage clamp recording showing photocurrent elicited by 1-second green light (530 nm) stimulation at 10 mw/mm^2^. (C) Quantification of the percentage of cells that showed action potential firing to a ramp current to 1x rheobase before (OFF) and during (ON) green light illumination. Optical illumination of bladder projecting Na_v_1.8-Arch neurons resulted in complete blockade of action potential firing at 1x rheobase. (D) Representative traces of action potentials elicited in Na_v_1.8-Arch-GFP-expressing neurons by ramp current injections of 1 to 4 times rheobase without (grey) and with (green) LED illumination. *** p=0.001. n=7 neurons; Paired t test. Error bars indicate SEM.

### Optogenetic inhibition of bladder sensory afferent terminals attenuates distension-induced pain and voiding behavior

We next tested if inhibition of Na_v_1.8^+^ bladder sensory terminals could attenuate bladder pain in response to distension. Bladder pain was assessed by the visceromotor response (VMR), evoked by bladder distention (10–60 mmHg distention in 10 mmHg steps, as we have described previously ^34,35^). Effects of light-dependent inhibition of Na_v_1.8^+^ afferents on bladder pain were tested by inserting a fiber optic cable via a “Y” tube that enabled delivery of compressed air to distend the bladder and light to illuminate the lumen of the bladder (Fig. 3A). In wild type mice, transurethral illumination with green light had no effect on the evoked VMR to bladder distention (Fig. 3B; D p>0.05. n= 12 mice per group; two-way ANOVA). However, in Na_v_1.8-Arch mice, transurethral illumination suppressed the distension-induced VMR compared to pre-illumination baseline VMR (Fig. 3C; E, p<0.05. n= 12 mice per group; two-way ANOVA). These results suggest that optogenetic inhibition of Na_v_1.8^+^ afferent terminals in the bladder can attenuate nociception evoked by bladder distension.

**Figure 3.**
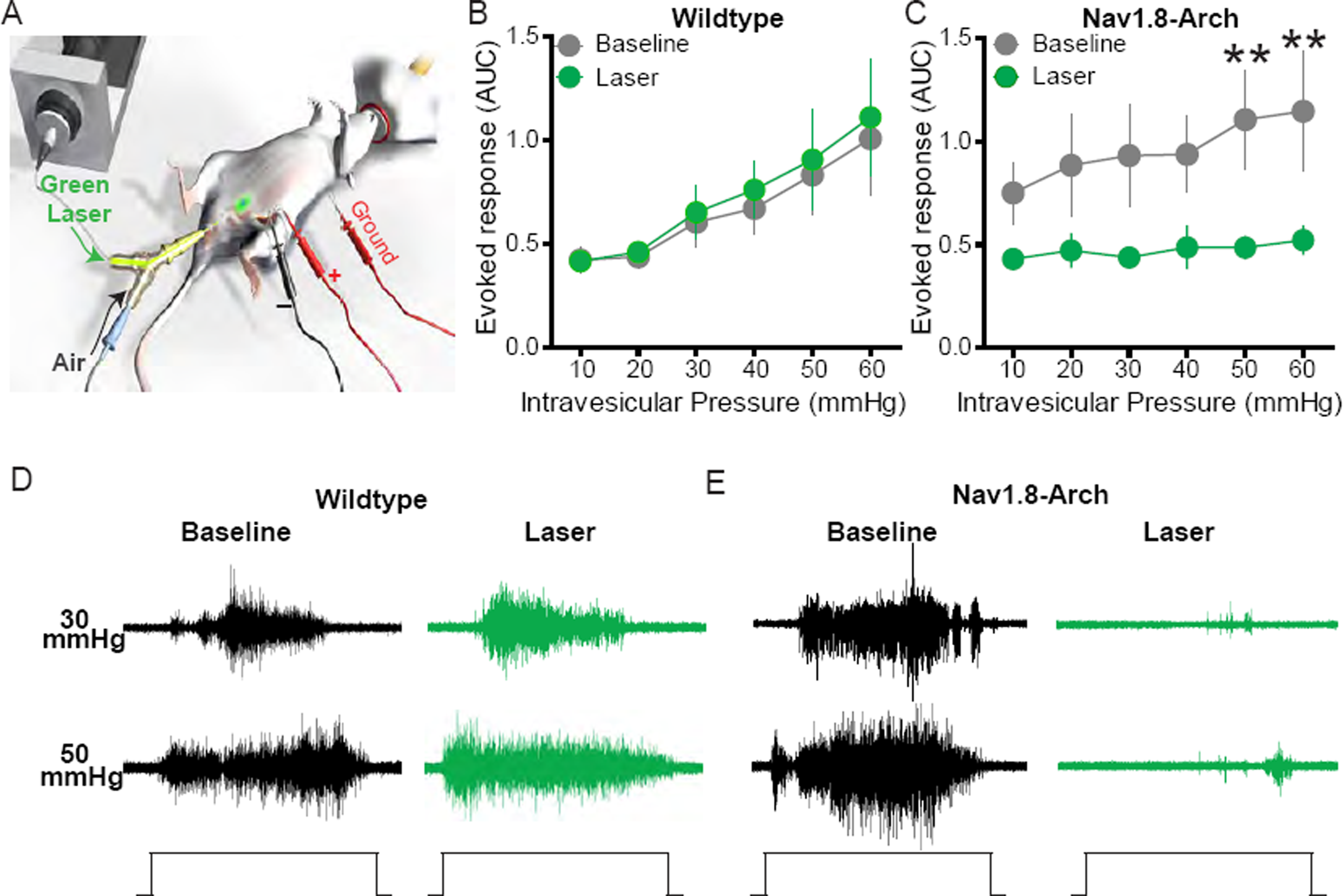
Optical silencing of Na_v_1.8^+^ bladder afferents attenuates bladder nociception. (A) Schematic illustrating bladder distention (VMR) setup to optically silence bladder afferents. (B & C) Transurethral fiber optic delivery of green light to the bladder lumen to induce optical silencing of bladder afferents significantly attenuated the evoked response to 50 and 60 mmHg bladder distention compared with baseline (pre-laser) responses in Na_v_1.8-Arch mice (** p<0.05. n= 12; F (1, 110) = 11.31), but had no effect in wild-type mice (p>0.05. n= 12; F (1, 110) = 0.7119). (D & E) Representative images of raw EMG traces from wild type and Na_v_1.8-Arch mice during 30 and 60 mmHg bladder distention taken before (baseline) and during (laser) green light illumination. ** p<0.05. n= 12 mice per group; F (1, 110) = 11.31, Two-way ANOVA. Error bars indicate SEM.

We determined if optogenetic inhibition of Na_v_1.8^+^ bladder afferents influences urodynamics by performing cystometric analysis of bladder function before and during laser illumination of the bladder (Fig. 4A). In anesthetized Na_v_1.8-Arch mice, illumination of bladder afferents with green light did not significantly alter maximum pressure, baseline pressure or threshold pressure between control or Na_v_1.8-Arch mice (Fig. 4 B, C). However, we did observe a slight (15.05%), yet statistically significant increase in the intercontraction interval (ICI) during laser illumination of the bladder in Na_v_1.8-Arch mice compared to control mice (Fig. 4 B-C; p=0.0183, n= 9-10 mice per group; Unpaired t test).

**Figure 4.**
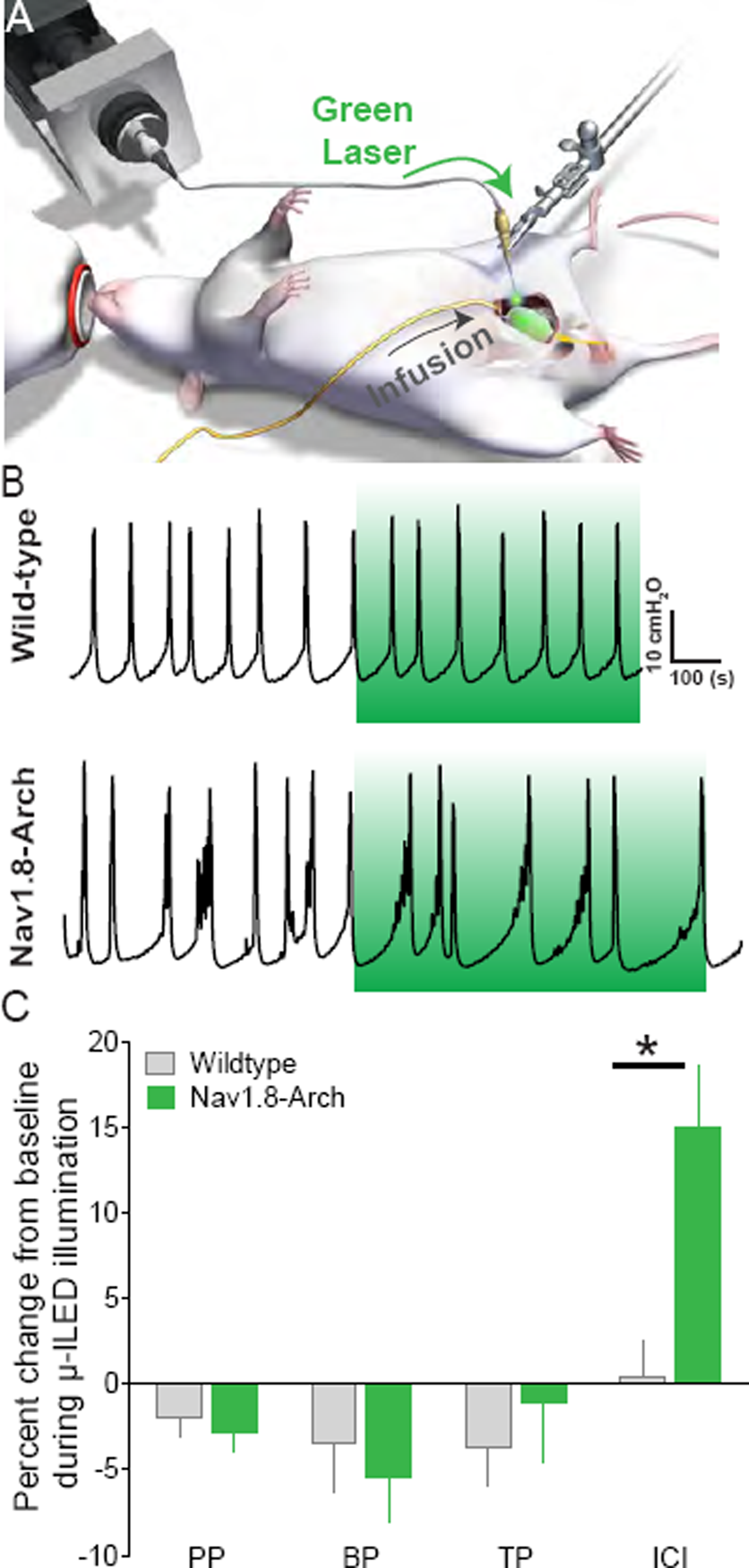
Effect of Arch induced inhibition of Na_v_1.8^+^ fibers on cystometric function. (A) Schematic illustrating cystometry setup using fiber optic to activate Arch in bladder sensory neurons. (B) Example cystometric traces recorded from wild type (top) and Na_v_1.8-Arch mice (bottom). (C) Quantification of peak pressure (PP), base pressure (BP), threshold pressure (TP) and intercontraction interval (IC) of wild type and Na_v_1.8-Arch mice before and during laser illumination of bladder normalized to baseline, comparing wild type to Na_v_1.8-Arch mice. Normalized ICI was significantly increased during laser illumination compared to wild type animals. * p=0.0183. n= 9-10 mice per group; Unpaired t test. Error bars indicate SEM.

### Development of a wireless optoelectronic device to enable optogenetic manipulation of bladder afferents in awake, freely moving animals

While the results above suggest a role for Na_v_1.8^+^ bladder afferents in mediating bladder pain and function, it is possible that the anesthetics used in these studies could interfere with pain processing. Currently, we lack technology to optically activate opsins expressed in the bladder afferents and determine their necessity in mediating bladder pain in awake, freely moving animals. We overcame this barrier by adapting technology that we have recently developed ^29,31^, to fabricate a flexible optoelectronic device to specifically target light to the lower abdomen. These devices are powered by near field communication (NFC) technology to achieve robust wireless functionality for chronic applications and for use in a wide variety of behavioral arenas (Fig. 5A). These wireless optoelectronic devices consist of a conductive metal pattern (18 μm thickness copper foil), semiconducting components (μ-ILED/mounted chips), with an encapsulation bilayer of polyisobutylene (PIB, 5 μm) and polydimethylsiloxane (PDMS, 500 μm). The electrical system incorporates a rectangular coil (width: 8 mm, length: 8 mm, copper traces: 7 turns with 70 um width and 30 um adjacent spacing) with surface mounted chips for power transfer via magnetic control to a loop antenna operating at 13.56 MHz. The overall dimensions of the device are 1 cm x 1 cm x 1 mm (*l* x *w* x *t*. Fig. 5B), which is depicted next to 200 mg capsules. An image of the device bent with forceps (Figure 5C) demonstrates the device flexibility and functionality in the bent conformation which enables interfacing with the abdominal wall. The thin-film PIB provides a protective barrier against moisture to allow for long-term use in a dynamic cage environment ^36^. This PIB barrier allowed for 100% of the devices to function for at least 4 weeks after implantation in the animal, with more than 50% of devices remaining functional after 6 months and some devices still functional at > 9 months after implantation.

**Figure 5.**
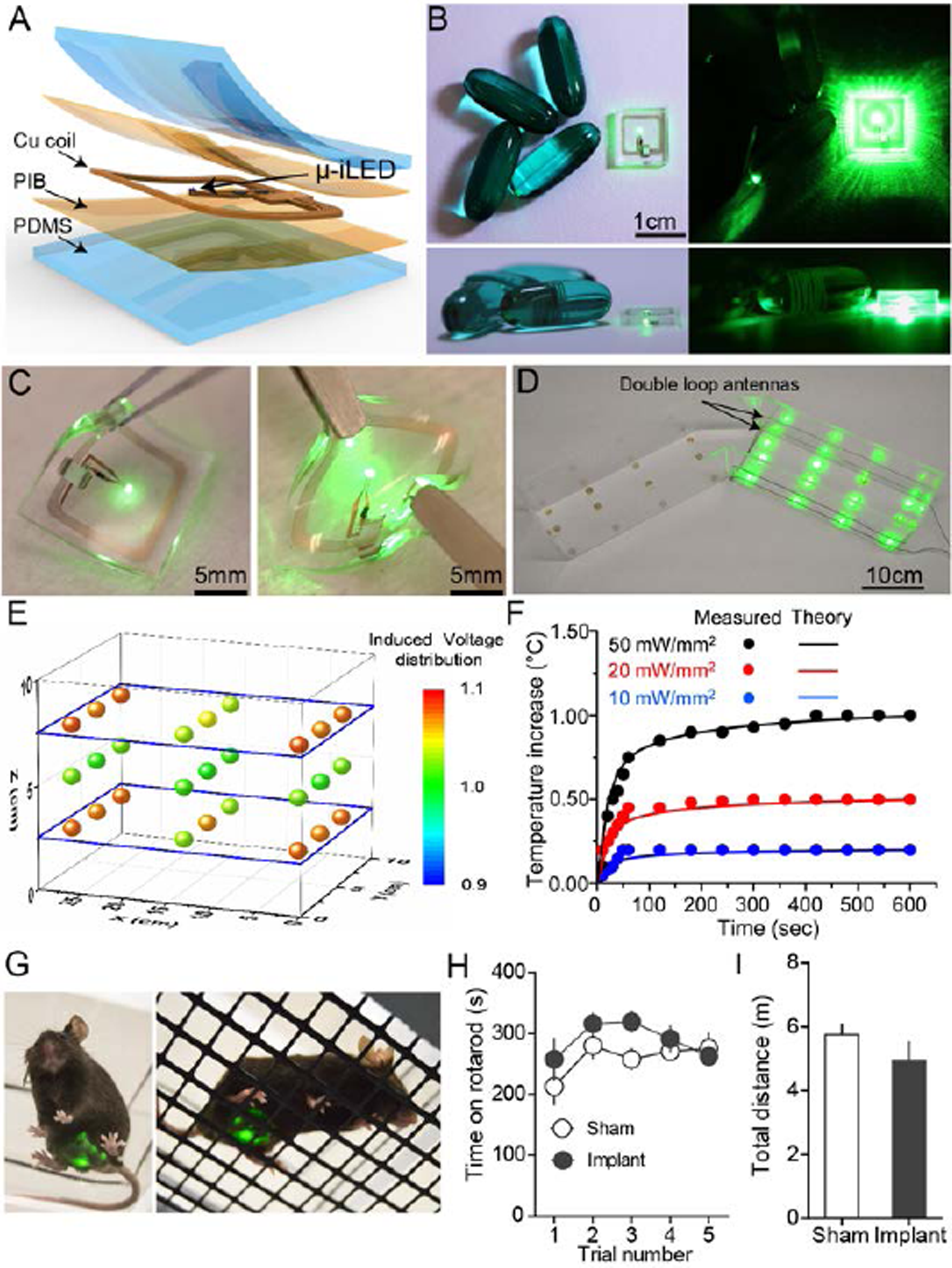
Flexible wireless optoelectronic device designed for optogenetic modulation of bladder afferent neurons. (A) Schematic illustration of the layered conformation of optoelectronic device. (B) Image of a micro-fabricated device adjacent to a 200 g capsules for the size comparison. (C) Demonstration of the flexibility of a functioning device with a forceps. (D) Image of wireless operation of optoelectronic devices in the V-maze with the double loop antennas. (E) Measurement of the normalized output power distribution of nine devices at heights of 2.5, 5, and 7.5 cm from the bottom of the cage demonstrating uniform power distribution of across the cage. (F) Thermal modeling and management of a μ-ILED device versus time at different peak output powers (20, 50, and 100 mW/mm^2^) demonstrating a maximum of 1 degree change at maximum light output of 100 mw/mm^2^. (G) Mice with wireless optoelectronic devices implanted over the bladder. (H) Implantation of the bladder optoelectronic device did not effect on motor behavior vs. sham animals in the rotarod test (P = 0.12, F_4,4_=1.118, t_1,8_=1.407 n = 8 sham, n = 8 device). (I) Mice with bladder implants did not exhibit significant difference in total distance travelled in open field test compared to sham mice (P = 0.16, two-tailed, n = 8 sham, n = 8 device).

These small and flexible optoelectronic devices can be adapted to a wide array of behavioral or home cage environments by tuning flexible signal antennas specifically for the dimensions of the behavioral or home cage environment. Figure 5D shows an example of the experimental setup for power transmission from antenna to optoelectronic devices in a V-maze (experimental data in Fig. 6B and C). The double-loop antenna is installed around the perimeter of one arm of the maze; for optimal power output performance, the loop antennae are placed 2.5 cm and 7.5 cm from the bottom of the cage. The spatial uniformity of power is demonstrated by equal illumination of twelve optoelectronic devices placed at different locations in a V-maze cage (Fig. 5D). This antenna configuration also allows for uniform μ-ILED illumination on the vertical dimension as well, which is important for maintaining consistent illumination during mouse rearing (Fig. 5E). The output power is normalized by its value at the center of double loop antenna at 5 cm and provides evidence that deviation of output power from the double loop antenna is less than 20%. This result indicates that the position of the double-loop antennas allows for a uniform magnetic field within the cage. This powering scheme allows for consistent and uniform illumination of implanted μ-ILEDs, which is critically important to reduce variability in light delivery during optogenetic experiments.

**Figure 6.**
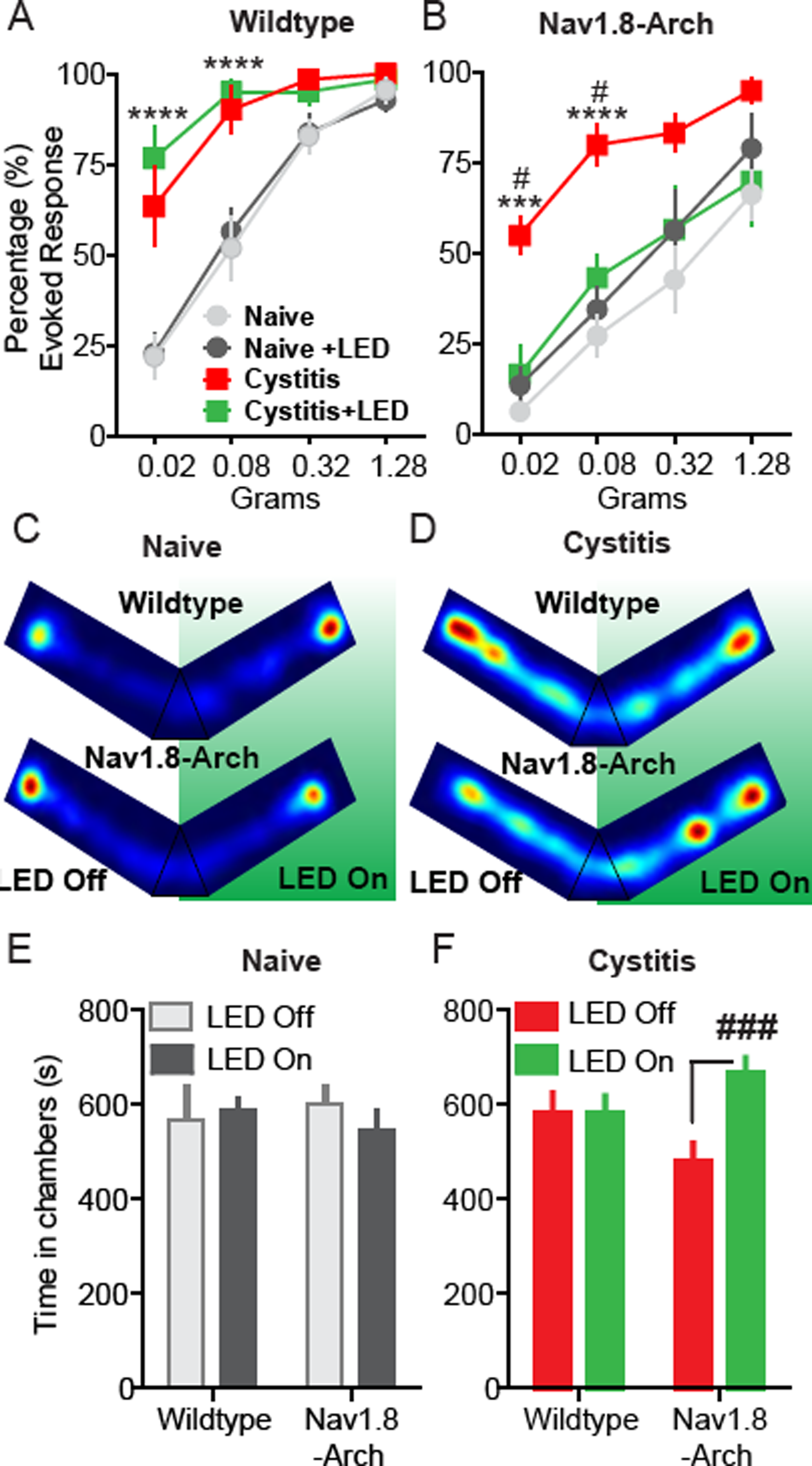
Optical silencing of Na_v_1.8^+^ bladder afferents attenuates evoked and ongoing bladder nociception. Quantification of abdominal mechanical sensitivity, measured by counting evoked responses to a series of von Frey filaments with increasing force before and during CYP-induced cystitis. (A) In wild type mice, activation of wireless μ-ILEDs had no significant effect on evoked responses in the naïve state compared to their baseline values. CYP (200 mg/kg i.p.) resulted in a significant increase in abdominal sensitivity. Activation of the optoelectronic devices implanted over bladder had no effect on abdominal sensitivity after cystitis (**** p<0.0001 comparing naïve to post-CYP, F (3, 90) = 76.31, n= 11 mice per group, two-way ANOVA). (B) In Na_v_1.8-Arch mice, activation of wireless μ-ILEDs had no significant effect on evoked responses in the naïve state compared to their baseline values. After cystitis induction, optical inhibition of Na_v_1.8^+^ sensory afferents significantly attenuated the abdominal hypersensitivity to values comparable to that of the baseline sensitivity recorded before CYP treatment (# p<0.05,*** p=0.0006, F (3, 30) = 7.596; **** p<0.0001, F (3, 90) = 67.17, n= 11 mice per group, two-way ANOVA). (C and D) Representative heat maps displaying time spent each zone of the custom V-maze in naïve (C) and cystitis (D) mice. Red indicates areas where the animals spend a higher proportion of their time. (E) In naïve conditions (pre-CYP), both wild type and Na_v_1.8-Arch mice did not exhibit any preference for either LED-ON or LED-OFF chamber (F_1,8_=0.21, p=0.6606, n= 6 mice per group, two-way ANOVA). (F) After cystitis induction, Na_v_1.8-Arch mice showed significant RTPP to the LED-ON arm compared to the LED-OFF arm, whereas wild type mice have no preference (F_1,18_=9.47, ### p=0.0065, n= 11 mice per group, two-way ANOVA). Error bars indicate SEM.

We also determined the heat generation under operation conditions by measuring the temperature at the surface of the device, in the mouse as a function of time, at peak output powers of 20 mW/mm^2^, 50 mW/mm^2^, and 100 mW/mm^2^. Measured values show good agreement with the thermal modeling, as shown in Figure 5F (symbols: measured values, lines; simulated theoretical values). The temperature approaches ‘steady-state’ after 5 min with a temperature increase of only 0.5 °C after 10 min at an output power of 50 mW/mm^2^. Mice with implanted NFC devices over the bladder exhibited no significant difference in locomotor function in the Rotarod test or in the open field test compared to sham-operated controls, suggesting that these devices do not impair motor behavior or coordination in freely moving animals (Fig. 5G-I).

### Wireless optogenetic inhibition of Na_v_1.8^+^ bladder sensory neurons reduces pain behaviors in a mouse model of bladder pain

Using the implantable optoelectronic devices described above, we evaluated the effects of optically inhibiting Na_v_1.8^+^ bladder afferents on referred abdominal sensitivity in awake, freely moving mice with CYP-induced cystitis. We have previously reported that patients with IC/BPS demonstrate referred hyperalgesia to the suprapubic region of the abdomen ^37^. Numerous studies have demonstrated that this referred hypersensitivity is reproduced in mouse models of bladder pain, including the CYP model ^34,37-44^. As it is not currently possible to induce bladder distension to measure VMR in awake (un-anesthetized) freely moving animals, we chose to test whether this referred abdominal hypersensitivity induced by CYP could be suppressed by inactivation of Na_v_1.8^+^ afferents. Na_v_1.8-Arch or wild type mice were implanted with wireless μ-ILED devices subcutaneously in the abdomen, with the μ-ILED positioned to illuminate the bladder through the abdominal musculature (Fig. 5G). After baseline assessments of cutaneous mechanical sensitivity, measured by application of von Frey filaments to the lower abdomen, mice were given a single injection of CYP (200 mg/kg, i.p), which induced robust referred mechanical hyperalgesia compared to baseline measurements (Fig. 6A-B) (pre-CYP, Fig. 6A, **** p<0.0001 comparing naïve to post-CYP, n= 11 mice per group, two-way ANOVA). Activation of the optoelectronic devices implanted over the bladder had no effect on abdominal sensitivity in either wild type or Na_v_1.8-Arch mice in the naïve state. However, optical inhibition of Na_v_1.8^+^ sensory afferents significantly reduced the abdominal hypersensitivity induced by CYP (# p<0.05, n= 6 mice per group, two-way ANOVA), an effect that is absent in wild type mice (Fig. 6A-B). The inhibition of Na_v_1.8^+^ sensory afferents in CYP-treated mice reduced abdominal mechanical sensitivity to values comparable to that of the baseline sensitivity (pre-CYP).

The results above demonstrate that referred mechanical hyperalgesia (von Frey filament experiments) can be reversed by optogenetic inhibition of Na_v_1.8^+^ bladder afferents. However, the primary complaint of IC/BPS patients is not referred hypersensitivity, but ongoing pain and pain associated with bladder filling ^37,45^. It is difficult to quantify ongoing pain in animals; in fact, it is not even clear if animals with CYP-induced cystitis exhibit ongoing pain. Development of these wireless optoelectronic devices presents us with the opportunity to evaluate ongoing, non-evoked visceral pain. We hypothesize that if silencing Na_v_1.8^+^ bladder afferents relieves ongoing bladder pain, then this should result in positive reinforcement (reward) on activation of the μ-ILED device. We can measure positive reinforcement using a real-time place preference (RTPP) assay, in which animals are placed in a V maze, with free access to roam throughout the maze. In one arm of the maze, NFC induces μ-ILED device activation (LED ON), while in the other arm, devices are inactive (LED OFF, Fig. 5D). Thus, in the LED ON arm, Na_v_1.8^+^ bladder afferents (which express Arch) are inhibited, and in the LED OFF arm, they can freely fire action potentials. If silencing Na_v_1.8^+^ bladder afferents relieves ongoing bladder pain, animals should demonstrate a preference for the LED ON arm compared to the LED OFF arm. In the naïve state, both wild type and Na_v_1.8^+^-Arch mice exhibited no preference for either the LED ON or LED OFF arms (Na_v_1.8-Arch LED ON vs LED OFF, p=0.6606, n= 6 mice per group, two-way ANOVA) (Fig. 6C; E). However, after CYP-induced cystitis, Na_v_1.8^+^-Arch mice showed significant RTPP for the LED ON arm compared to the LED OFF arm, whereas wild type mice with CYP-induced cystitis show no preference (Na_v_1.8-Arch LED ON vs LED OFF, ### p=0.0065, n= 11 mice per group, two-way ANOVA) (Fig. 6D-F). These results suggest that inhibition of Na_v_1.8^+^ bladder sensory afferents attenuates both ongoing pain and referred abdominal hypersensitivity in a mouse model of bladder pain.

## Discussion

The cardinal clinical symptoms of IC/BPS include pain upon bladder filling (distention) and inhibiting bladder sensory afferents can reduce this pain in many patients. Prior studies have demonstrated that the majority of bladder-projecting C-fiber neurons express Na_v_1.8-mediated Na^2+^ currents ^20,46^. In addition, C-fiber bladder afferents that express Na_v_1.8 channels are known to exhibit increased excitability in animal models of bladder pain, suggesting the importance of these afferents in mediating bladder nociception ^23,24^. We confirmed the assumptions made by these original studies ^23,24^ by specifically inhibiting Na_v_1.8^+^ bladder afferents with the inhibitory channel Arch expressed under the control of the Na_v_1.8 promoter and demonstrated that these fibers play a critical role in bladder nociception and a minor role in bladder voiding.

Previous studies have shown that C-fibers are silent during normal bladder distension and can become active during bladder injury. Thus, C-fiber activity is thought to contribute to bladder nociception in response to injury and have a sparse role in innocuous sensation of the bladder under normal physiological conditions ^20,46-49^. Consistent with these reports, we find that inhibition of Na_v_1.8^+^ bladder sensory afferents significantly attenuates nociception related to bladder injury induced by CYP and has no effect in naïve mice.

Somewhat surprisingly, in our cystometry studies, we found that silencing Na_v_1.8^+^ bladder sensory afferents resulted in a small yet significant increase in the ICI in naïve Na_v_1.8-Arch mice. The first possible explanation for this is that the relatively non-physiological, rapid rate of bladder filling in continuous flow cystometry engages nociceptive afferents to a small degree and inhibition of these afferents leads to delays in ICI. Alternatively, the effect of silencing Na_v_1.8^+^ afferents on bladder function may not be due to Na_v_1.8 expression in small diameter nociceptors. Indeed, consistent with a prior report ^50^, we demonstrate that a proportion of Na_v_1.8^+^ bladder-projecting neurons co-express NF200 (42 ± 10.9%), a marker of large myelinated sensory neurons. It is possible that these cells include low threshold mechanoreceptors, which are known to be involved in the voiding reflex ^51,52^. Thus, this population of bladder afferents contributes to the delay in ICI we observe after optically silencing Nav1.8^+^ bladder afferents.

A limitation of using VMR to measure bladder nociception is that these experiments must be performed in lightly anesthetized mice, and the response is evoked with invasive bladder distention. The fully implantable wireless NFC-powered μ-ILED devices we developed offer the ability to optically control neural activity in awake (un-anesthetized) and freely behaving mice. This allows us to determine the effects of silencing Na_v_1.8^+^ bladder sensory afferents in both normal and pathological states. Optical silencing of Na_v_1.8^+^ bladder afferents resulted in attenuation of referred mechanical hypersensitivity induced by CYP, while there were no effects of green light in naïve mice. This result suggests that in the context of bladder pain, hyperactivity in the Na_v_1.8^+^ afferents mediates referred mechanical hypersensitivity. One limitation of this experiment is that we cannot be certain whether silencing Na_v_1.8^+^ bladder afferents or Na_v_1.8^+^ cutaneous afferents resulted in this attenuation of referred mechanical hypersensitivity, as light from our devices could also inhibit cutaneous abdominal afferents over the device. Nonetheless, these studies demonstrate that Na_v_1.8^+^ afferents are critical for the mechanical hypersensitivity evoked by the CYP model of bladder pain.

While IC/BPS patients demonstrate referred hypersensitivity, the primary clinical complaint is spontaneous pain and pain on bladder filling ^37,45,53^. Reflexive measurements of referred hypersensitivity do not assess whether the ongoing bladder pain induced by cystitis is being relieved. It is possible, for example, that silencing Na_v_1.8^+^ bladder afferents attenuates mechanical hypersensitivity but does not reduce ongoing pain associated with bladder filling. We employed a RTPP paradigm to evaluate whether silencing the Na_v_1.8^+^ bladder afferents causes relief of spontaneous or ongoing bladder pain. Inhibition of Na_v_1.8^+^ bladder afferents produced robust RTPP in mice with cystitis, but not in uninjured mice without any cystitis. This result suggests that CYP-induced cystitis indeed induces ongoing pain, and that relief of this ongoing pain is analgesic and rewarding as evident by robust RTPP. To our knowledge this is the first demonstration of optogenetic inhibition of primary afferent neurons reducing spontaneous or ongoing bladder pain, or pain of any origin.

Combining cell type specific optogenetic inhibitory opsins with fully implantable wireless μ-ILEDs offers insights into the potential application of optogenetic neuromodulation to the treatment of pathological pain. To achieve clinical translatability of optogenetics, several obstacles remain, including the development of viral gene therapy vectors that are safe and effective for use in humans. It is vital to develop a robust vector that can efficiently target a large number of bladder afferents, but at the same time achieve cell type-specific targeting, as this could help reduce unwanted side effects. However, assessment of potential toxicities associated with long-term expression of opsins in neurons will be critical. Promising advances have been made in gene therapy by targeting neurons using viral vectors, but much work must be done to advance this to the clinical arena. The recent advent of precise gene editing technologies could allow the delivery of opsins to specific subsets of afferent neurons innervating the bladder ^54,55^. The bladder is easily accessible to these innovative approaches. Combining any of these gene therapy approaches that prove to be safe with real-time control of neural dynamics with closed-loop optoelectronic systems ^56-59^ could lead to the development of future therapies for bladder pain and/or voiding dysfunction. The wireless devices we have developed could be easily scaled for use in humans, and our results suggest that optogenetic silencing of Na_v_1.8^+^ bladder afferents could be an effective approach to alleviate pain associated with IC/BPS.

## Materials and Methods

### Animals and genetic strategy

Adult mice (8–12 weeks of age) were used for this study, unless otherwise noted. Mice were housed in the animal facilities of the Washington University School of Medicine on a 12 h light/dark cycle, with ad libitum access to food and water. Institutionally approved protocols were followed for all aspects of this study. Experiments were performed on male heterozygous Na_v_1.8-cre mice^32^ crossed to female homozygous Gt(ROSA)26Sor^tm35.1(CAG-aop3/GFP)Hze^ (Ai35) mice from Jackson Laboratory carrying floxed stop-Arch-GFP gene in the Gt(ROSA)26Sor locus^33^ to generate the Na_v_1.8-Arch mouse line.

### Immunohistochemistry

Immunohistochemistry was performed using the same methods as Park et al ^29^. Briefly, DRG tissue was collected 7 days after Cholera Toxin Subunit B Alexa Fluor^®^ 555 (Thermo Fischer Scientific, C22843) injection. Tissues were fixed with 4% PFA and then embedded with O.C.T. Compound (Tissue-Tek, 4583) for sectioning. Tissues were washed in PBS and incubated in blocking solution (10% normal goat serum/0.1% Triton-X/1x PBS) for 1 hour at room temperature. Primary antibodies (1:200 Mouse anti-CGRP, Sigma C7113; 1:200 Mouse anti-NF200, Sigma N0142; 1:1000 Rabbit anti-GFP, Thermo Fischer A11122), or IB4 (1:100, IB4 Alexa Fluor 647, Thermo Fischer I32450) were diluted in blocking solution and incubated on the sections overnight. Slides were washed 3x for 10min each with PBS and incubated with secondary antibodies diluted in blocking solution for 1 hour at room temperature (1:1000 Goat anti-mouse IgG Alexa Fluor 647, Thermo Fischer A-21235; 1:1000 Goat anti-rabbit IgG Alexa Fluor 488, Thermo Fischer A11008). Slides were washed 3x for 10min each with PBS, and allowed to dry before mounting coverslips (Vectashield Hard Set, H-1400). Samples were imaged using a Leica TCS SP5 confocal microscope. For more detailed methods see supplementary methods.

### Dorsal root ganglion (DRG) culture and whole-cell electrophysiology

DRG culture and whole cell recordings were performed as in Park et al ^29^. Briefly, DRG neurons were dissociated from mice 7 days after DiI (ThermoFisher) injection. Neurons were recorded and optically stimulated with an EPC10 amplifier (HEKA Instruments) and Patchmaster software (HEKA Instruments). Optical stimulation was delivered through the microscope objective, using a custom set-up with a green (530nm) LED (M530L3; Thorlabs). Light intensity of the LED at the focal plane was 10 mW/mm^2^. For more detailed methods see supplementary methods.

### Visceromotor reflex behavior

The visceromotor reflex (VMR) in female mice was quantified using abdominal electromyograph (EMG) responses. The VMR is a reliable behavioral index of visceral nociception in rodents and was performed as previously described ^60-64^. Briefly, mice were anesthetized with isoflurane (2% in oxygen) and silver wire electrodes were placed in the oblique abdominal muscle. A lubricated, 24-gauge angiocatheter was passed into the bladder via the urethra for urinary bladder distension (UBD). After surgical preparation, isoflurane was reduced to ∼1% until a flexion reflex response was present (evoked by pinching the paw), but righting reflex was absent. Phasic UBD consisted of graded distensions at pressures of 10-60 mmHg. Baseline EMG activity was subtracted from EMG during UBD, rectified, and integrated to obtain distension-evoked EMG responses. Distension-evoked EMG is presented as area under the curve. Experimenter was blinded to mouse genotype. For more detailed methods see supplementary methods.

### Cystometry

Mice (8-22 weeks old) were first anesthetized with isoflurane (2%), then a midline incision was used to expose the bladder dome. A 25-gauge butterfly needle was used to puncture the dome of the bladder and warm mineral oil placed on the exposed bladder to prevent tissue drying. After the catheter was placed, anesthesia was lowered to 1.5% for 1 hour. Then over 30-40 minutes, the anesthesia was slowly reduced to ~0.8%. At this point, bladders were filled at 0.04 ml/min with room-temperature water or saline (no differences observed between fluids; data not shown) using a syringe pump to evoke a regular voiding pattern. Intravesicular pressure was measured using a pressure transducer amplified by a Transbridge transducer amplifier (WPI) and recorded using WINDAQ data acquisition software (DataQ Instruments) at a sampling rate of 5 Hz. After a regular voiding pattern was established, a ten-minute baseline was collected followed by laser illumination (described below) for at least ten minutes and a ten-minute recovery period. The laser was always turned on at the end of a contraction and then off at the end of a contraction resulting in some trials that were greater than ten-minutes. These trials were completed in duplicate for each animal and parameters were averaged. Data were analyzed using a Matlab (Mathworks) script to determine base pressure (BP), threshold pressure (TP), maximum pressure (MP) and intercontraction interval (ICI) (terminology conformed to ^65^). All data files were blinded so analysis could occur in an unbiased manner.

### Photoinhibition of VMR and Cystometry

Optical inhibition of VMR was performed using a 532 nm, 200 mW diode-pumped solid-state (DPSS) laser. In visceromotor reflex studies, a fiber optic (200 μm diameter core; BFH48-200-Multimode, NA 0.48; Thor labs) was coupled to the laser and connected to the transurethral catheter via a Y-shaped connector. The fiber tip was positioned 0.1 mm beyond the tip of the catheter in the bladder lumen. Photoinhibition was performed at 20 mW/mm^2^ constant illumination. For cystometry, photoinhibition was performed with the same fiber optic positioned above the exposed bladder dome instead of transurethrally and at 30 mW/mm^2^ constant illumination.

### NFC device fabrication

Micro-fabrication of the soft, flexible wireless optoelectronic devices started with a spin-cast poly(dimethylsiloxane) (PDMS, Sylgard 184) at 600 RPM x 60 second on a clean glass slide (75 x 50 x 1 mm, L x W x thickness), followed by curing at 150°C x 10 min. Next, an 8% by weight solution of polyisobutylene (PIB, BASF) in heptane was applied to freshly cured PDMS by spin-casting at 1000 RPM x 60 second followed by a 3 min bake at 100°C. Then copper foil (18 μm thickness) was placed on the newly formed polymer substrate. Photolithography (AZ 4620, AZ Electronic Materials) and copper etching was then used to define the conducting pattern. Next, the electronic components (μ-ILED (TR2227, 540 nm, Cree Inc. Raleigh, NC), rectifier (CBDQR0130L-HF, Comchip Technology, Freemont, CA), and 136 pF capacitor (GRM1555C1H680JA01J, Murata electronics, Japan)) were placed onto the copper pattern using conductive paste. Then a second layer of polyisobutylene was added to the device, followed by another PDMS layer, using the same conditions described above.

### Implantation of wireless NFC device

The NFC optoelectronic devices were used to activate Arch in freely moving mice. Under isoflurane anesthesia, a small incision was made in the abdominal skin and the device was implanted subcutaneously between the skin and muscle, positioned to illuminate the lower abdomen and bladder. The incision was then closed using surgical staples. The animals were allowed to recover for at least 4 days before behavioral experiments were performed.

### Cyclophosphamide (CYP) induced cystitis

Cyclophosphamide (CYP) was used to induce bladder inflammation and visceral pain. Bladder cystitis was initiated by a single injection of CYP (200 mg/kg; i.p; Sigma, St. Louis) dissolved in saline. Behavioral assays were performed before (baseline), 4 and 24 hours after CYP injection.

### Abdominal mechanical sensitivity

Abdominal sensitivity was measured by counting the number of withdrawal responses to 10 applications of von Frey filaments (North Coast Medical, Inc, Gilroy, CA; 0.02, 0.08, 0.32 and 1.28 g) to the lower abdomen. Each mouse was allowed at least 15 seconds between each application and at least 5 minutes between each size filament. Animals were acclimated to individual boxes on a plastic screen mesh for at least one hour before testing. All experimenters were blinded to mouse genotype and treatment. NFC devices were activated by an antenna wrapped around the individual cage, prior to the experiment the antenna was tested to insure full NFC coverage of the cage. NFC antenna power and signal were generated by Neurolux hardware and software (Neurolux, Urbana, IL).

### Real-time place preference (RTPP)

Place preference was tested in a custom made V-maze constructed of plexiglass with a layer of corn cob bedding. Each arm of the two-arm V maze is 10 cm wide × 30 cm long x 10 cm height neutral area between arms. To generate the NFC signal, one arm of the maze was covered with an NFC-emitting antenna (Neurolux, Urbana, IL) allowing for the control of μ-ILED devices throughout one arm of the maze. To begin the experimental protocol, a mouse was placed in the neutral area of the maze and was continuously monitored and recorded through a video connection for 20 min. Ethovision software (Noldus, Leesburg, VA.) was used to determine time-in-chamber and generate representative heat maps for each condition.

### Statistics

Results are expressed as means ± SEM. Mann-Whiney test was used to compare the percentage suppression of action potentials and cytometric parameters. To analyze VMR and mechanical sensitivity data, two-way ANOVA with repeated measures was used. Bonferroni’s *post hoc* tests were used (when significant main effects were found) to compare effects of variables (genotype, treatment). A value of *p* <0.05 was considered statistically significant for all statistical comparisons. Researchers were blinded to all experimental conditions. Two replicate measurements were performed and averaged in all behavioral assays.

## Data Availability

The datasets generated during and/or analysed during the current study are available from the corresponding author on reasonable request.

## Acknowledgments

This work was funded by the NIH Director’s Transformative Research Award (TR01 NS081707) and an NIH SPARC award (NIBIB U18EB021793) to RWG and JAR, and R01NS42595 to RWG, Urology Care Foundation Research Scholars Program and Kailash Kedia Research Scholar award to VKS. McDonnell Center for Cellular and Molecular Neurobiology Postdoctoral Fellowship to A.D.M.

## Author Contributions

V.K.S., A.D.M., and R.W.G. conceptualized the project; V.K.S. and A.D.M. designed, performed experiments and collected data. J.G.G. performed anatomical analysis. M.P. performed electrophysiological experiments. G.B.G. performed behavioral experiments. S.V. performed mouse genotyping. H.H.L provided equipment for the VMR and cystometry experiments. K.N.N. provided Matlab code for cystometry data analysis. J.Y., K.C. and J.A.R. designed and provided optoelectronic devices. V.K.S., A.D.M. and R.W.G. analyzed the data and wrote the manuscript with comments from all the authors.

